# Challenges in Detecting Somatic Recombination of Repeat Elements: Insights from Short and Long Read Datasets

**DOI:** 10.1101/2024.08.25.609631

**Authors:** Giovanni Pascarella, Martin Frith, Piero Carninci

**Affiliations:** RIKEN Center for Integrative Medical Sciences (IMS), Yokohama 230-0045, Japan; Artificial Intelligence Research Center, National Institute of Advanced Industrial Science and Technology (AIST), Tokyo 135-0064, Japan; Graduate School of Frontier Sciences, University of Tokyo, Chiba 277-8562, Japan; Computational Bio Big-Data Open Innovation Laboratory (CBBD-OIL), AIST, Tokyo 135-0064, Japan; Human Technopole, Milan 20157, Italy

## Abstract

Non-allelic copies of the two major families of repeat elements in the human genome, Alu and L1, recombine somatically at high frequency. Tissue-specific recombination profiles are dynamic in cell differentiation and are altered in neurodegeneration, suggesting that somatic recombination of repeat elements can contribute to functional heterogeneity of cells in health and disease. The study of these genomic variants, however, presents several technical challenges related to their extremely low copy number and their sequence content. Here, we address key issues regarding detecting and annotating structural variants derived from recombining repeat elements in NGS data. We show that PCR introduces significant changes of recombination profiles in sequencing libraries and that recombination profiles are affected by the choice of sequencing platform. We refine previous estimates of recombination in single cells by analyzing recombination profiles in PCR-free HG002 datasets sequenced by Oxford Nanopore Technologies and PacBio sequencers while describing several platform-specific differences. We additionally provide evidence that recombination events annotated in state-of-the-art single-cell HG002 whole-genome sequencing datasets are likely molecular artifacts generated by PCR. By exploring the limits of current technologies, this work establishes essential requirements for future developments to enhance the reliability of detecting somatic recombination of repeat elements in genomic datasets.

## Introduction

The genomes of different individuals are distinguished by sequence variants that are either inherited or generated in the germline. Variants can also arise later in development and be confined to specific tissues, cell types, or even single cells, contributing to somatic diversity and disease (Poduri et al. 2013; Horebeek et al. 2019; Olafsson and Anderson 2021; Yu et al. 2024). Variants are broadly classified by size as short sequence variants (<50bp) and structural variants (“SV”, >50bp). While annotating all variants in an individual is currently unattainable, focusing on somatic variants prevalent in specific tissues or cell types may reveal susceptibility to genomic instability and illuminate disease processes (Poduri et al. 2013; Vijg and Dong 2020). However, studying somatic variants is challenging due to their rarity, often supported by only a few sequencing reads, necessitating highly sensitive and specific workflows. Cross-validation is often impossible because genomic copies may be depleted during library preparation. As a result, variants supported by sparse reads are frequently overlooked in genomic studies. We recently reported that non-allelic homologous repeat element sequences recombine in somatic cells, generating deletions, inversions, duplications, and translocations, detected at unexpectedly high frequencies using a novel bioinformatic pipeline (Pascarella et al. 2022). Most of our findings were based on short-read libraries enriched for Alu and L1 repeat sequences (“capture-seq”), in which we estimated up to 5 NAHR events per cell. Although we demonstrated that genome-wide profiles or recombination events supported by just one split read differ significantly from random recombination generated *in silico*, our estimates may be influenced by technical biases, including tissue sampling bias, uneven capture probe efficiency, loss of sensitivity due to short reads, PCR amplification bias, and false positives from chimeric amplicons generated by PCR. These last two points are of particular relevance in protocols focused on repeat sequences, since PCR can produce artifacts resembling SVs (Pääbo et al. 1990; Meyerhans et al. 1990; Ji et al. 1994; Kurahashi et al. 2000; Kanagawa 2003). In this work, we initially sought to verify whether PCR can introduce artifacts resembling NAHR of repeat elements in capture-seq libraries by performing our capture-seq protocol on selected genomic regions encompassing a small set of human Alu repeats. We show that chimeric amplicons with the same sequence features of NAHR events can be generated even at low PCR cycling. Prompted by these results, we performed a multi-platform, comprehensive assessment of the reproducibility of NAHR detection in short and long-read datasets generated by international consortia for human genome assembly and benchmarking of SV annotation. We report several sequencing platform-specific differences in genomic profiles of NAHR and discuss possible underlying reasons. Finally, we test the detection limits of NAHR in state-of-the-art single-cell datasets to show that existing protocols are not yet suitable for studying NAHR of repeat elements in single cells.

## Results

To understand whether PCR can generate chimeric artifacts that resemble NAHR of repeat elements in NGS libraries, we performed our capture-seq protocol on a pool of known input genomic fragments encompassing Alu sequences. We designed forward and reverse primer pairs on genomic regions flanking 16 AluS and 4 AluY (Figure 1A and Supplemental Table S1).

**Figure 1.**
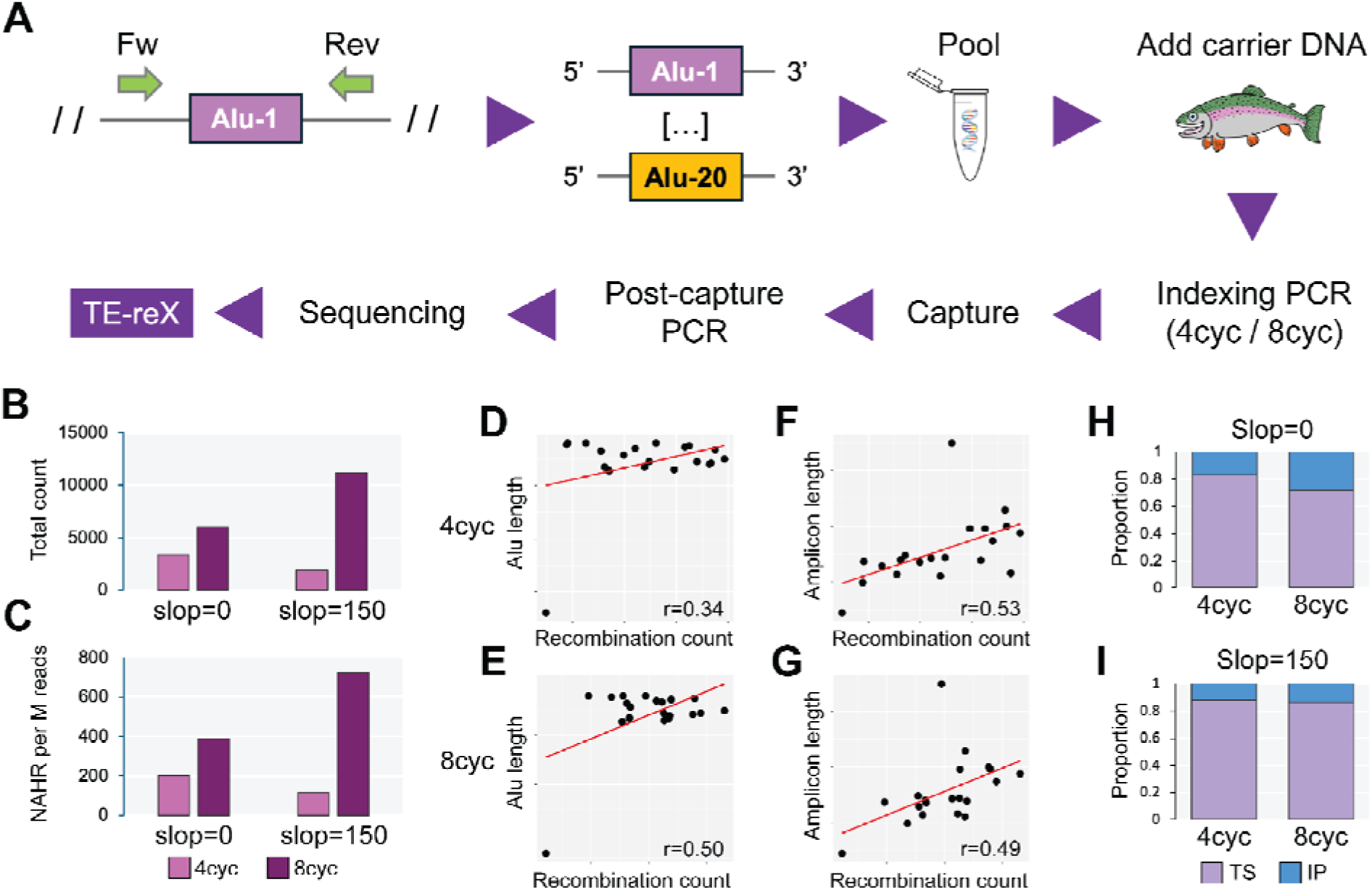
Identification of chimeric NAHR-like molecular artifacts in test capture-seq libraries. **A)** Schematics of the generation, sequencing, and analysis of the test capture-seq libraries. **B, C)** Raw and normalized counts of chimeric recombination artifacts in 4cyc and 8cyc libraries detected by TE-reX with slop=0 and slop=150. **D-G)** Dowplots showing the correlation between the count of chimeric recombination events per input Alu sequence with its respective length or the length of the amplicon, including each Alu plus the non-repeat 5’ and 3’ flanking sequences in 4cyc and 8cyc libraries with slop=150. **h, i)** Relative proportion of chimeric recombination events generated by template switching (“TS”) or internal priming (“IP”) in 4cyc and 8cyc libraries as detected by TE-reX run with slop=0 (h) and slop=150 (i).

**Supplemental Table S1.**
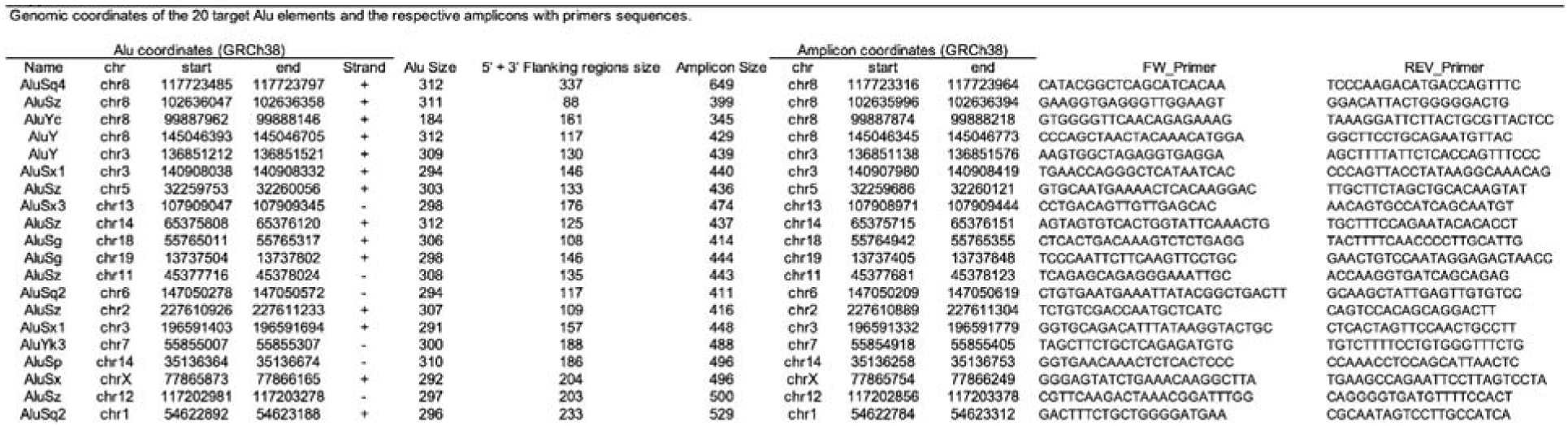
Genomic coordinates of the 20 target Alu elements and the respective amplicons with primers sequences.

The pairwise sequence identity rate across all targets was 81%±4% (Supplemental Figure S1 and Supplemental Table S2).

**Supplemental Figure S1.**
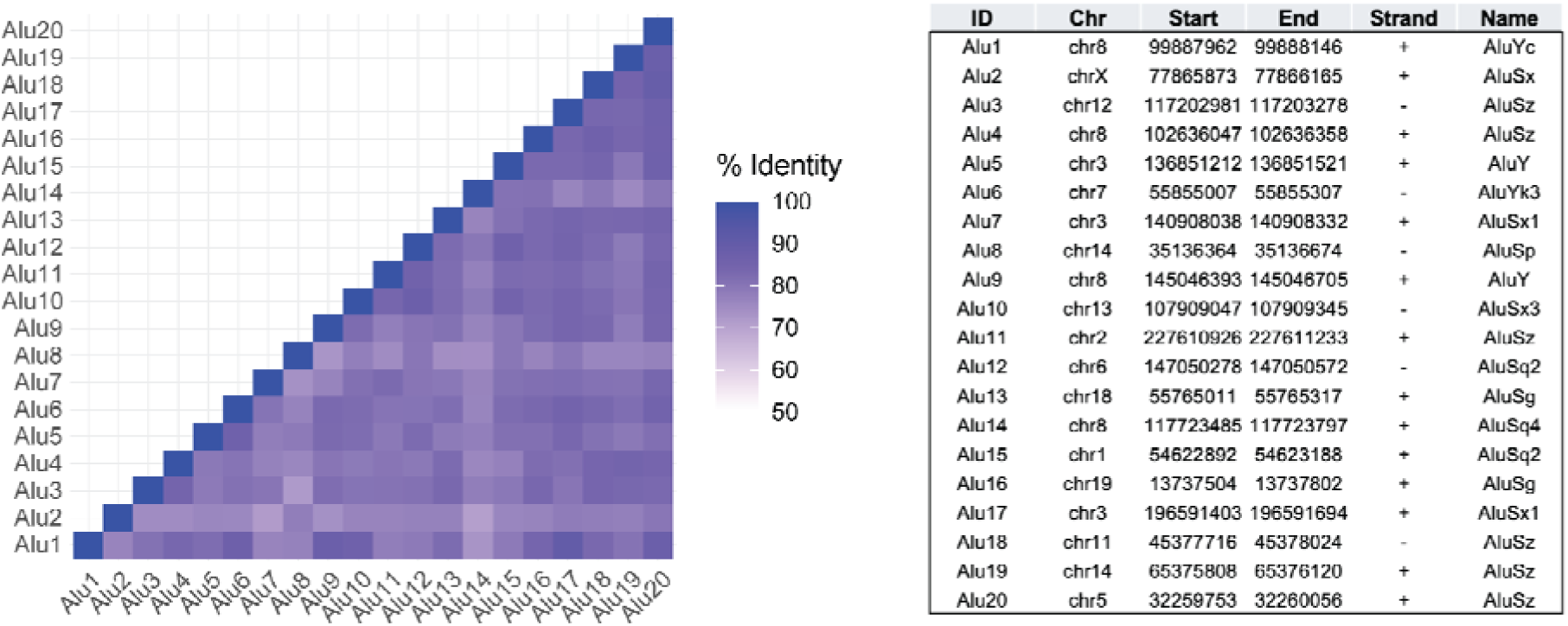
Pairwise matrix of identity rate for the 20 Alu elements (“seq1-20”) included in the test capture pool. The table on the right shows the identity of the 20 Alu elements in the left plot.

**Supplemental Table S2.**
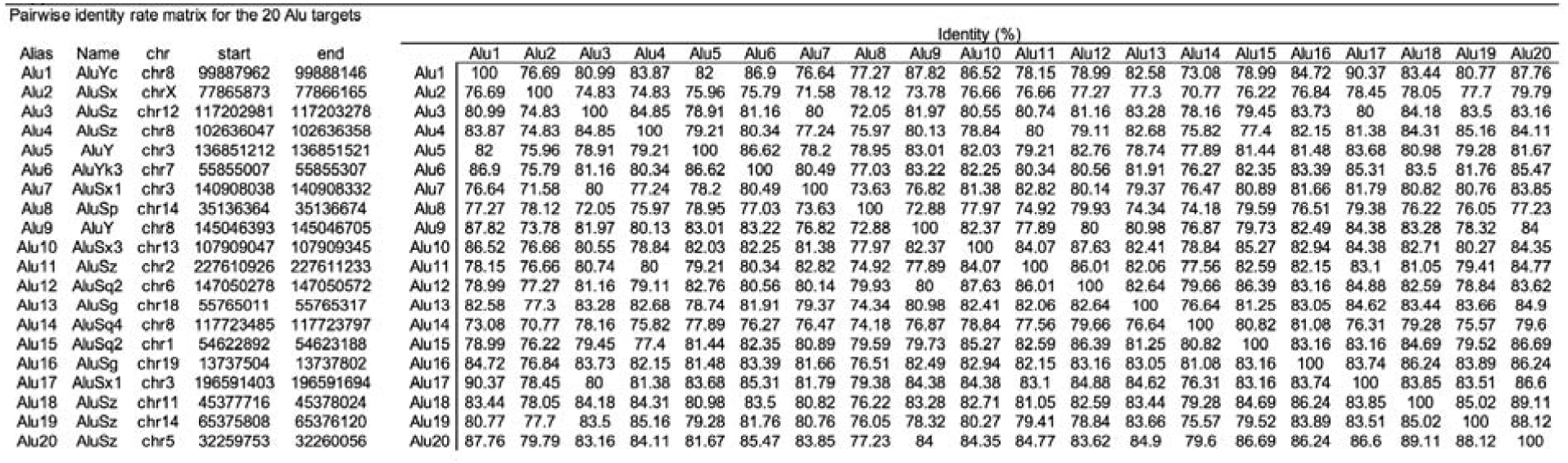
Pairwise identity rate matrix for the 20 Alu targets.

The average size of the amplicons was 457bp ± 62bp, allowing end-to-end coverage in the sequenced reads. We performed PCR from a genomic DNA sample obtained from the amygdala of a neurotypical donor, and we verified the correct amplification of the 20 fragments by agarose gel and Sanger sequencing (Supplemental Figure S2).

**Supplemental Figure S2.**
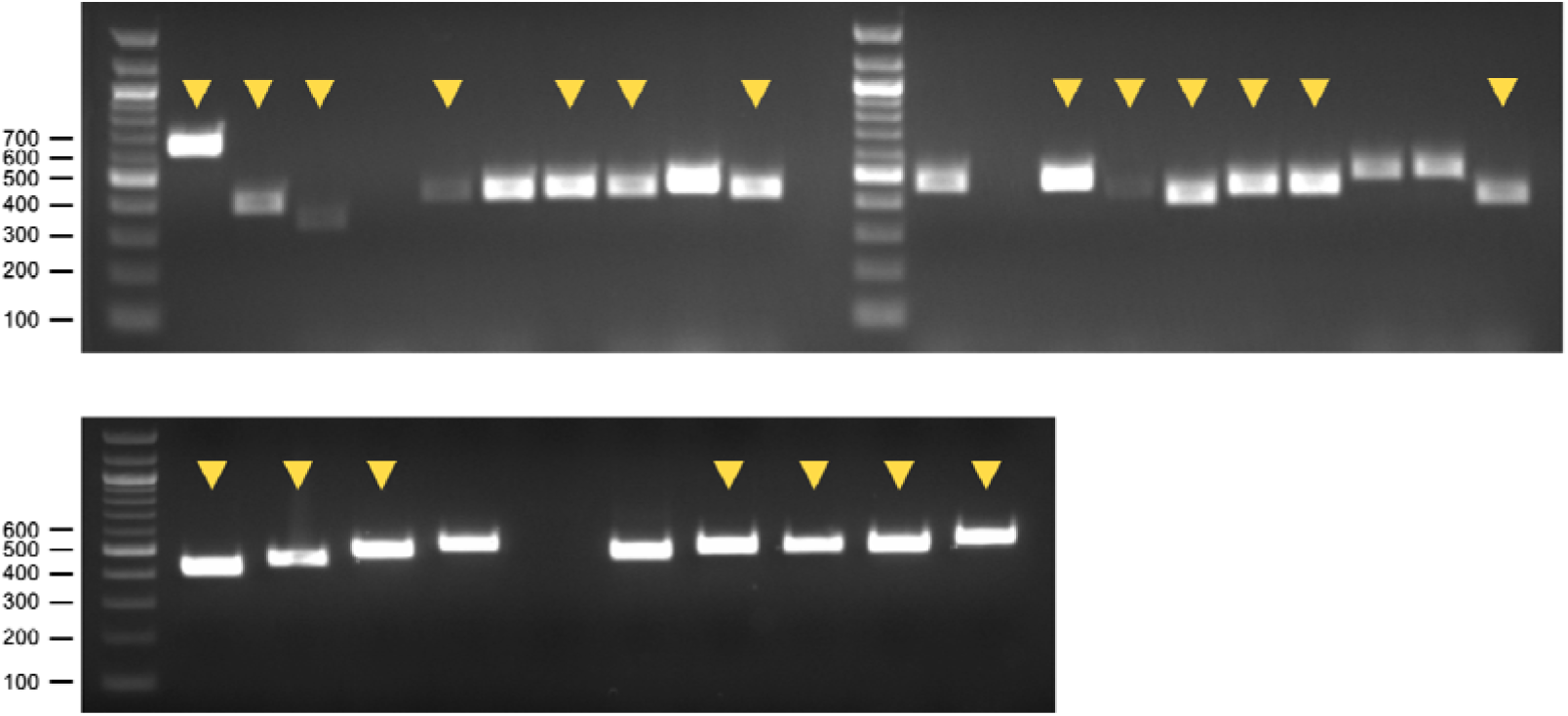
Gel electrophoresis of the 20 amplicons included in the capture pool (yellow arrowheads).

We pooled the 20 amplicons in an equimolar fashion; to obtain a sample replicating the proportion of Alu sequences in the human genome (∼10%), we mixed the pooled amplicons in a 1:10 ratio with fragmented Salmon genomic DNA, an inert carrier commonly added to hybridization reactions. To test if chimeric recombination can be introduced by the PCR generally required to introduce library indexes, we generated two indexed pre-capture libraries that were amplified for 4 PCR cycles (“4cyc”) or 8 cycles (“8cyc”). We enriched the targets using our capture-seq protocol (Pascarella et al. 2023); the captured fragments were then amplified with 12 PCR cycles, pooled equimolarly and sequenced by Illumina Miseq, generating respectively 17 million reads (4cyc) and 15 million reads (8cyc). After mapping the reads to GRCh38, we used the TE-reX pipeline to find potential chimeric recombination events. TE-reX annotates NAHR of repeat elements from reads that, after genome alignment, are split on different loci containing homologous repeat elements. A key parameter of TE-reX is “slop”, which indicates the offset of the two genomic breakpoints of a split read from positions of perfect homology, as defined by repeat model sequences in the Dfam database. By setting slop to 0 (“-s 0” or “--slop=0” in TE-reX command line), TE-reX annotates NAHR events; values higher than 0 return recombination events with breakpoints at non-homologous positions that may result, for example, from non-homologous end joining (NHEJ) (Supplemental Figure S3).

**Supplemental Figure S3.**
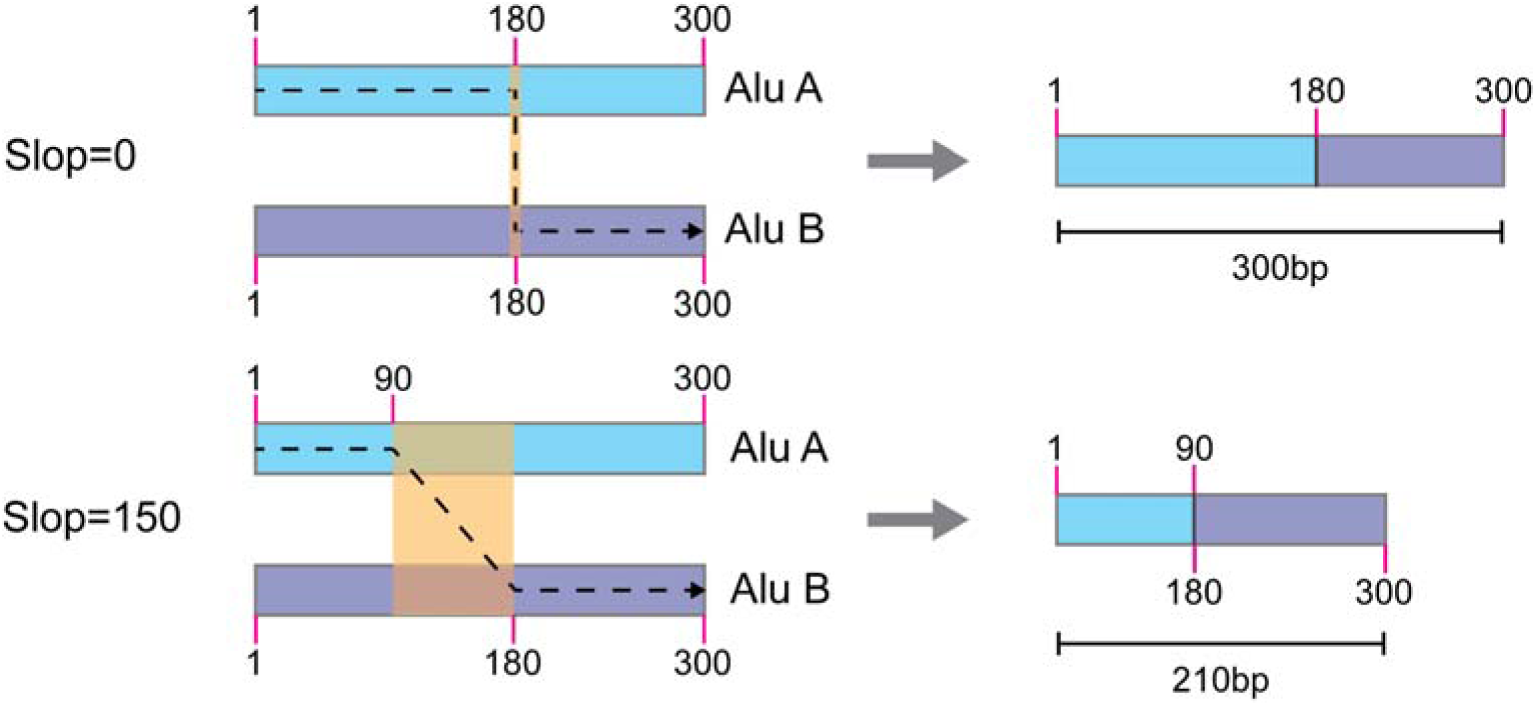
Effect of different slop settings in TE-reX. With slop=0. TE-reX annotates recombination events where repeats are joined at homologous positions, determined from the repeat models deposited in the Dfam database. Recombination events annotated with slop>0 feature repeats joined at non-homologous positions. In the example in the figure, TE-reX with slop=0 identified a non-allelic recombination event with repeats Alu A and Alu B joined at the homologous position 180 on both split reads. The resulting hybrid Alu has the same genomic size as the original repeats. With slop=150, TE-reX identified a recombination event where Alu A and Alu B are joined at position 90 and 180, respectively. The resulting hybrid Alu is shorter than each of the 2 original repeats, since it contains a deletion (region shaded in orange color).

TE-reX detected chimeric recombination of the 20 input sequences in both 4cyc and 8cyc libraries. With slop=0, the total number of chimeric events was 3442 in 4cyc and 5869 in 8cyc; a normalization by the sequencing depth returned 202 chimeric recombination events per million sequenced reads in 4cyc and 379 in 8cyc (Figure 1B-C). The near 2-fold difference between 4cyc and 8cyc confirmed that the pre-capture indexing PCR could already generate chimeric artifactual recombination of Alu elements. We extended the search for chimeric rearrangements that do not follow the criteria of perfect homology at breakpoints by running TE-reX with slop=150 (a value that equals half of a model Alu sequence, 300bp); TE-reX returned 1915 Alu-Alu rearrangements in 4cyc and 11204 in 8cyc (respectively 112 and 726 events per million sequenced reads, Figure 1B-C). We detected all possible rearrangement permutations of the 20 input Alu elements (n=190) in both 4cyc and 8cyc with slop=0, while with slop=150, we found 184 combinations in 4cyc and all 190 in 8cyc (Supplemental Table 3).

**Supplemental Table S3.**
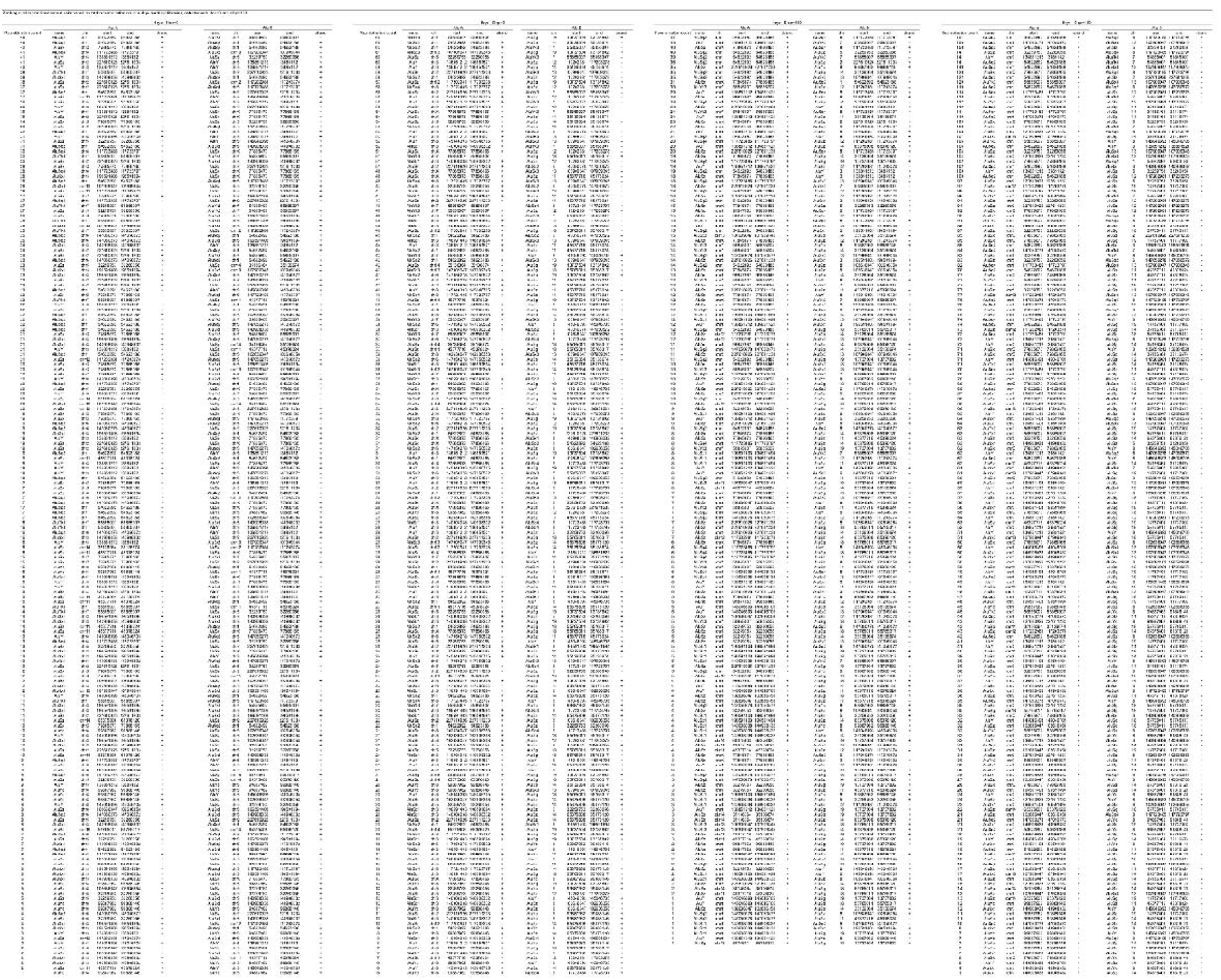
Ranking of all recombined repeat pairs based on total recombination count in 4cyc and 8cyc libraries, detected with slop=0 and slop=150.

By ranking the number of chimeric recombinations detected per Alu element, we found that some Alu sequences were more likely to recombine than others (Supplemental Table S4). We reasoned that sequence identity rate would be the most likely factor driving Alu-Alu recombination. However, we did not detect any correlation between the identity rate of recombined Alu pairs and their respective artifacts count (Supplemental Figure S4).

**Supplemental Table S4.**
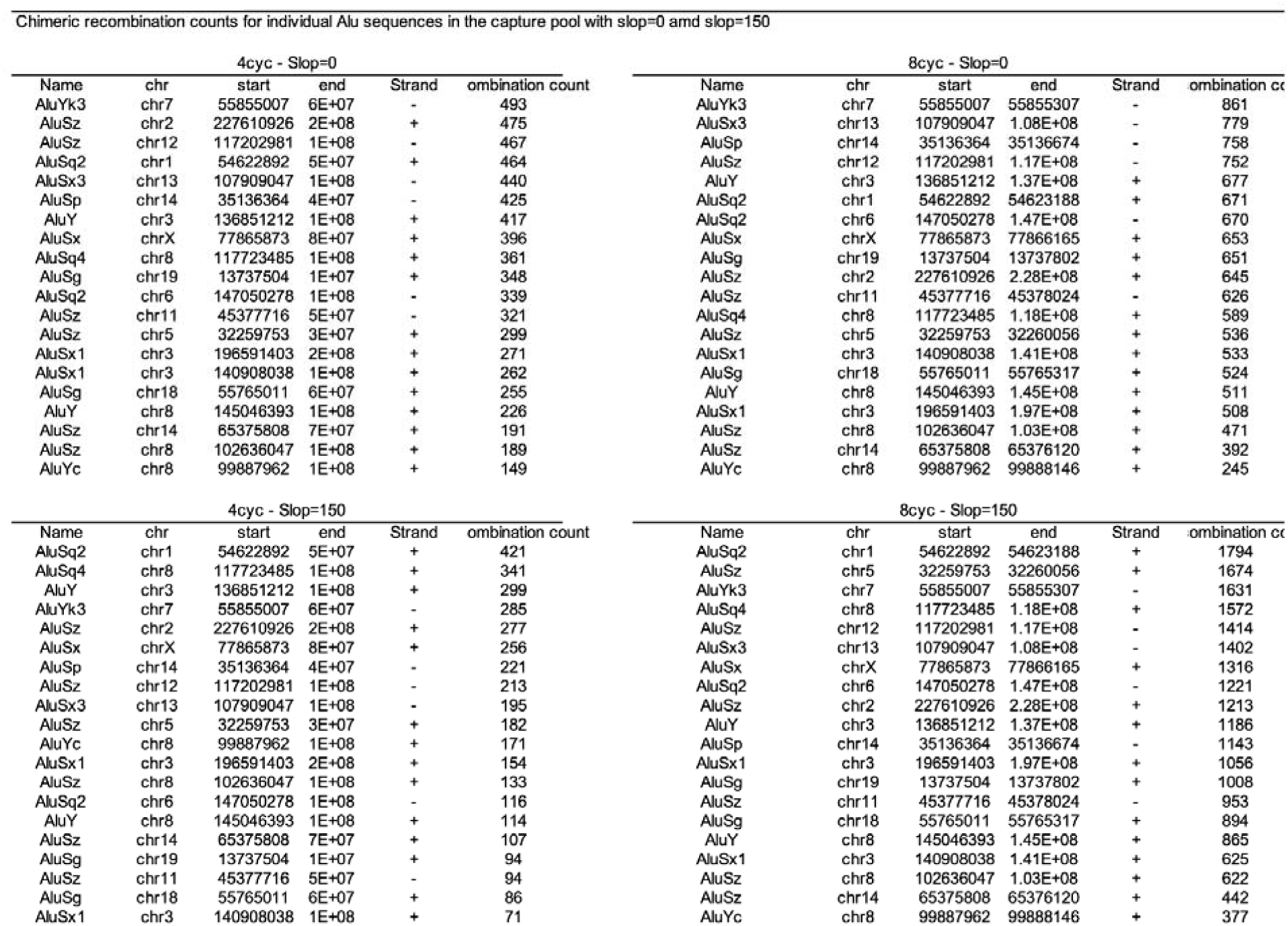
Chimeric recombination counts for individual Alu sequences in the capture pool.

**Supplemental Figure S4.**
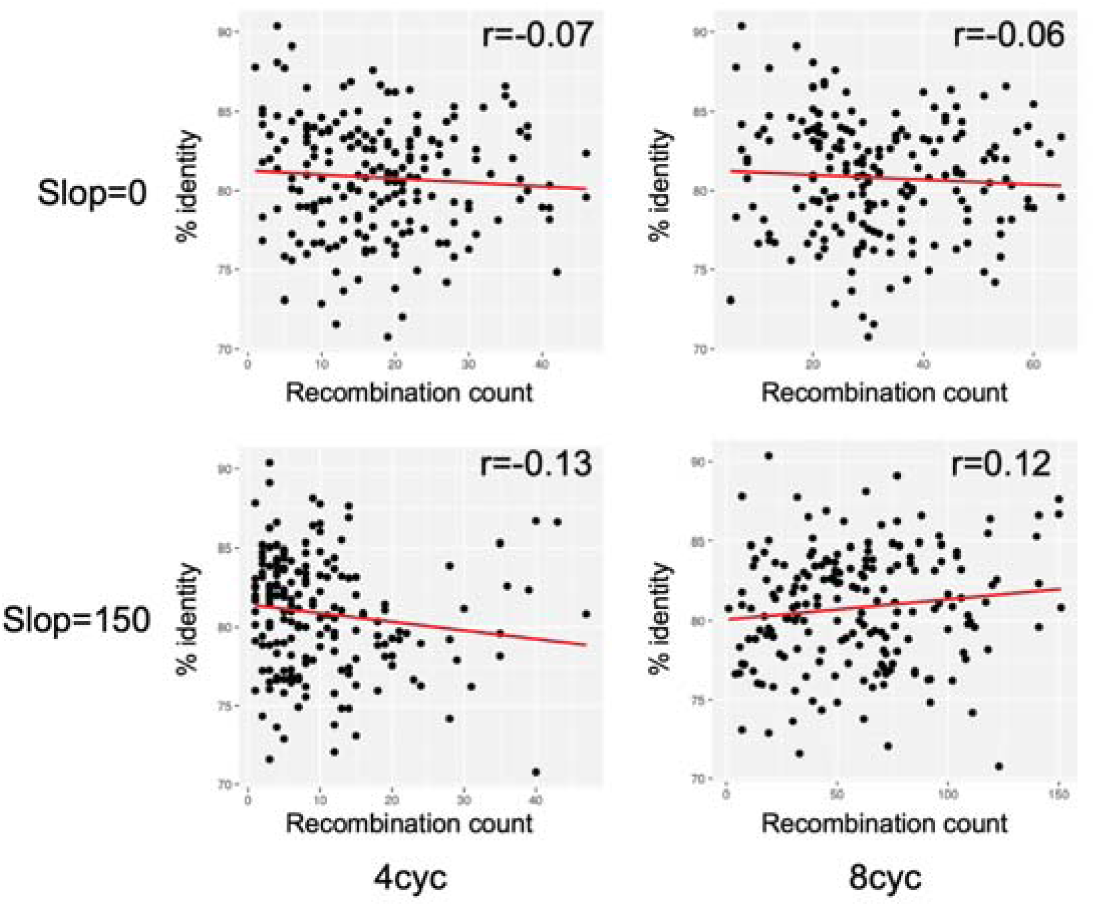
Dotplots showing the correlation between the count of chimeric recombination events per Alu pairs with the respective sequence identity rates in 4cyc and 8cyc libraries with slop=0 and slop=150.

Extending our analyses to other sequence features, we found that the number of chimera per Alu elements was better correlated with the total length of each amplicon than with the length of the encompassed Alu repeats, even though the 5’ and 3’ genomic sequences flanking the Alu elements shared no homology with each other (Figure 1D-G and Supplemental Figure S5).

**Supplemental Figure S5.**
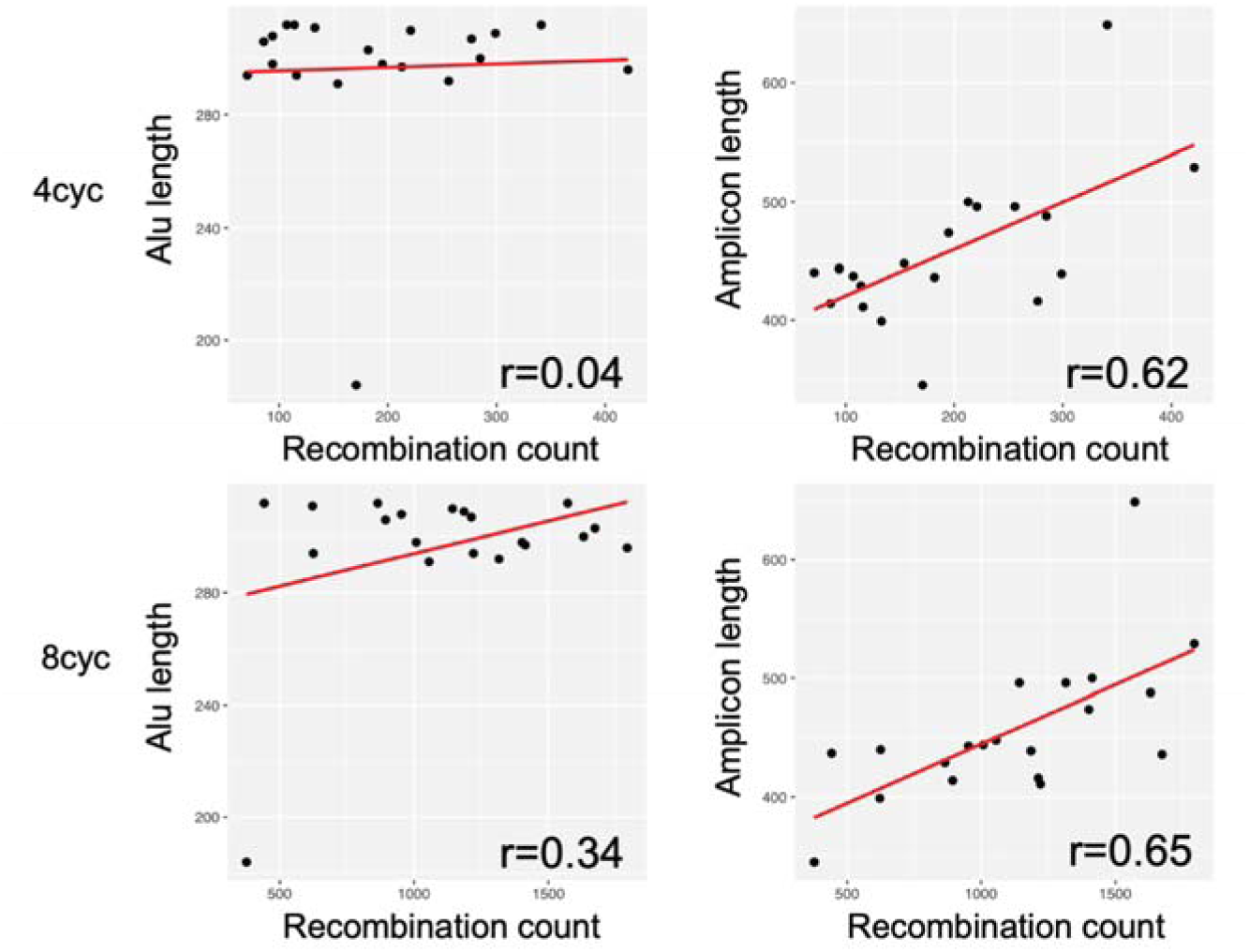
Dotplots showing the correlation between the count of chimeric recombination events per input Alu sequence with its respective length or the length of the amplicon including each Alu plus the non-repeat 5’ and 3’ flanking sequences in 4cyc and 8cyc libraries with slop=150.

This suggested that the total template length is essential to confer stability during the formation of chimeric molecules. The main mechanisms that can lead to the generation of recombination artifacts are template switching (“TS’’) and internal priming (“IP”). TS occurs when the DNA polymerase “jumps’’ on an alternative template that presents sequence identity to the template that is undergoing amplification. Chimeric amplicons formed by TS should comprise the full-length 5’ to 3’ sequence, with a sequence switch present at some point along the homology region; this sequence architecture makes artifacts formed by TS indistinguishable from NAHR events. On the other hand, IP artifacts originate when one amplified molecule with homology to other molecules in the template pool acts as a primer and triggers an illegal extension. As a result, IP artifacts have 5’ or 3’ truncations with respect to the original template sequences. We analyzed the sequences of the artifacts detected in the 4cyc and 8cyc libraries to identify split reads without (TS) or with sequence truncations (IP). We found that both with slop=0 and slop=150, a large majority of recombination artifacts were generated by TS, since they retained their 5’ and 3’ repeat-flanking regions (Figure 1H-I). Overall, these results illustrate that, in principle, NGS libraries that rely on PCR may contain chimeric molecules with architectural features resembling NAHR, which could potentially have consequences for downstream analyses. However, the genome of each cell is characterized by a much more complex reservoir of repeat sequences than the pool used in these test capture-seq libraries; consequently, the occurrence of chimeric NAHR-like artifacts in real-world libraries may differ. To verify this, we obtained a dataset originally aimed at benchmarking various genomics analytical methods using the samples of the Genome in a Bottle consortium (GIAB) (Baid et al. 2020; data from gs://brain-genomics-public/research/sequencing/fastq). The GIAB samples consist of lymphoblastoid cell lines generated from 7 donors (HG001 to HG007, of which HG002-4 and HG005-7 are two son-mother-father trios of Ashkenazi and Han descendants; Zook et al. 2014, 2019). The dataset consists of 40x whole-genome sequencing libraries generated without (“PCR-”) or with PCR (“PCR+”, n=5 cycles), and sequenced by two high-throughput NGS Illumina sequencers (Hiseq X Ten and Novaseq6000). After aligning the libraries on GRCh38 and annotating NAHR by TE-reX, when comparing PCR-vs PCR+ libraries, or Novaseq vs Hiseq datasets, we did not detect any significant difference in the number of NAHR events normalized by sequencing depth or in estimates of the number of NAHR per diploid cell, (Figure 2A).

**Figure 2.**
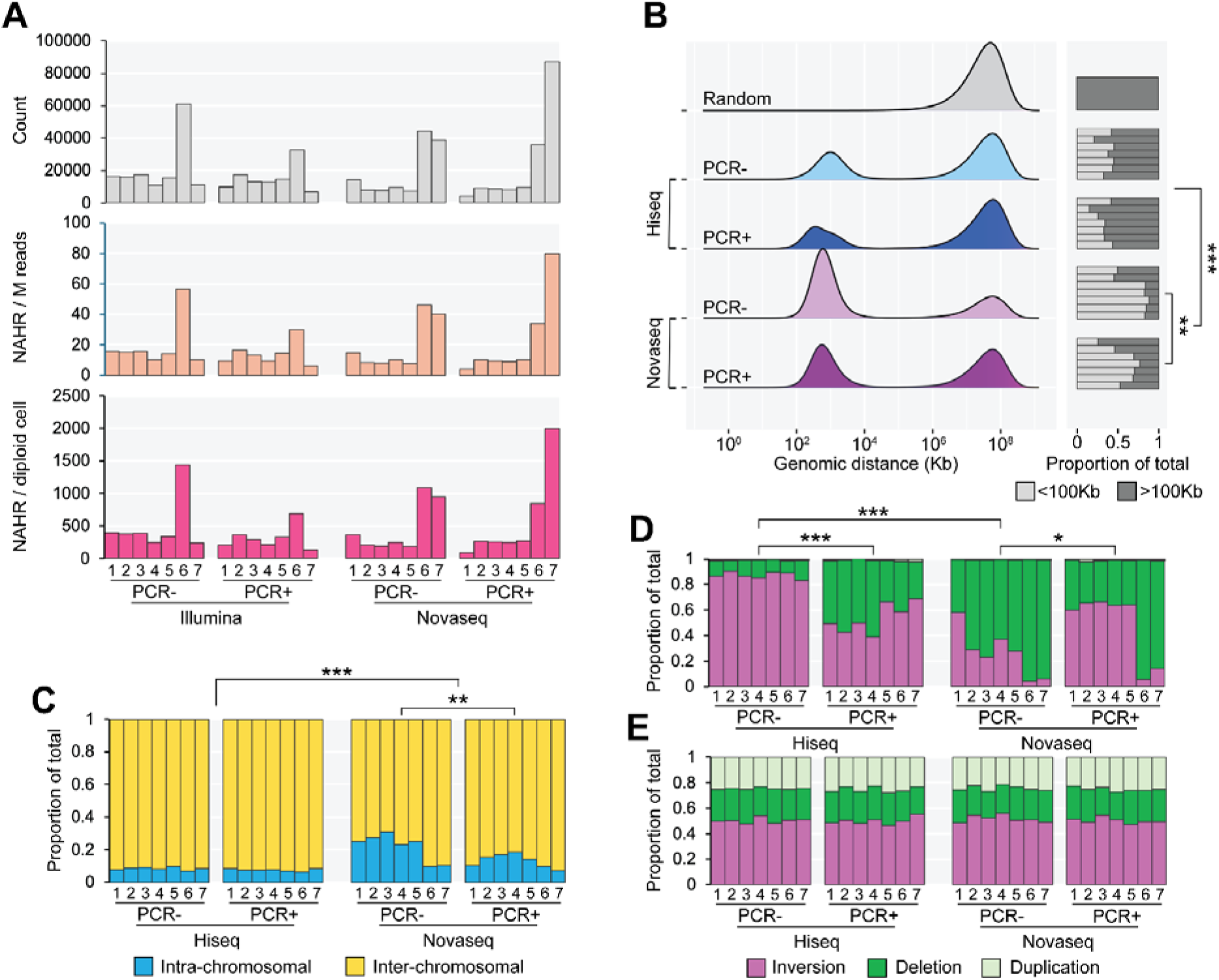
Analysis of NAHR in a WGS dataset of GIAB samples reveals significant cross-platform and PCR-induced differences. **A)** Counts of NAHR in the WGS dataset of GIAB samples (top: raw counts; middle: counts normalized by millions of reads per library; bottom: estimated NAHR events per diploid cell). **B)** Relative proportion of intra- and inter-chromosomal NAHR. **C)** Genomic distance of repeat pairs involved in intra-chromosomal NAHR. The barplots on the right-hand side show the relative proportion of recombined proximal (d<100Kb) and distal (d>100Kb) repeat pairs. **D, E)** Relative proportion of SV types generated by NAHR, for proximal (D) and distal (E) repeat pairs. ^∗^p < 0.05; ^∗∗^p 0.01; ^∗∗∗^p < 0.001 (Student’s T-test). For the bar plots, numbers at the bottom represent the respective library names (HG001-HG007). Each bar represents the results for a single library.

However, a comparison of the genome-wide profiles of NAHR revealed several notable differences. PCR+ Novaseq libraries showed a higher rate of inter-chromosomal NAHR than the PCR-libraries, and the Novaseq dataset had overall higher rates of intra-chromosomal recombination compared to the Hiseq dataset (Figure 2B). We observed a similar pattern when looking at the genomic distance of repeat pairs recombined within the same chromosome; Novaseq PCR-libraries had a higher proportion of NAHR between repeat elements distanced less than 100Kb (“proximal”) compared to the PCR+ libraries, and the Novaseq dataset had more recombination of proximal repeats than the Hiseq dataset (Figure 2C). Intra-chromosomal NAHR of repeats in inverted configuration generates an inversion of the intervening genomic region, while NAHR of repeats in direct configuration can generate a single deletion, or a coupled duplication and deletion (Sasaki et al. 2010). In PCR-free conditions, the Novaseq dataset displayed more deletions than the Hiseq dataset. The PCR increased the proportion of deletions in the Hiseq dataset, whereas the opposite was true for the Novaseq dataset, where the PCR increased the proportion of inversions (Figure 2D). These differences were detected only for proximal NAHR events, with distal NAHR generating similar profiles of SVs in the two datasets (Figure 2E). Overall, these results indicate that profiles of NAHR of repeat elements can be affected by the choice of different short-reads high-throughput sequencing platforms. Moreover, although it does not affect the overall counts and estimates of NAHR per cell, including even a few PCR cycles in library generation can significantly alter the intra-chromosomal profiles of NAHR.

These discrepancies in NAHR profiles justified further characterization of NAHR in libraries sequenced using alternative methodologies and platforms. Of all the GIAB samples, the lymphoblastoid cell line HG002, generated from the son of the Ashkenazi trio, has been used in recent years for genome assembly and variants annotation by the Telomere-to-Telomere Consortium (T2T) and the Human Pangenome Reference Consortium (HPRC) (Nurk et al. 2022; Jarvis et al. 2022). As a result of the scientific endeavors from these consortia, the HG002 genome has been extensively sequenced at deep coverage with instruments producing long DNA reads from Oxford Nanopore Technologies (“ONT”) and Pacific Biosciences (“PacBio”). We obtained the fastq files for 22 PCR-free WGS libraries sequenced on ONT PromethION, and 22 PCR-free libraries sequenced on PacBio HiFi Revio (https://s3-us-west-2.amazonaws.com/human-pangenomics/index.html?prefix=T2T/scratch/HG002/sequencing/ont/) (https://s3-us-west-2.amazonaws.com/human-pangenomics/index.html?prefix=T2T/scratch/HG002/sequencing/hifi/) (https://s3-us-west-2.amazonaws.com/human-pangenomics/index.html?prefix=NHGRI_UCSC_panel/HG002/hpp_HG002_NA24385_son_v1/PacBio_HiFi/).

The main advantage of these datasets is the availability of multiple high-coverage technical replicates of a single cell type obtained from a single donor performed from homogeneous DNA samples.

For the ONT dataset, all libraries were prepared from a single DNA pellet obtained from large cultures of HG002 (personal communication from Miten Jain). While technical notes available for the PacBio dataset report similar conditions (https://github.com/human-pangenomics/HG002_Data_Freeze_v1.0?tab=readme-ov-file) (https://s3-us-west-2.amazonaws.com/human-pangenomics/NHGRI_UCSC_panel/HG002/hpp_HG002_NA24385_son_v1/PacBio_HiFi/PacBio_HiFi_HG002_README.txt), we were not able to directly confirm this, or to verify whether the PacBio libraries were produced from DNA of the same HG002 batch as the ONT dataset. Despite these limitations, we considered these datasets as a valuable resource to investigate the reproducibility and sensitivity of detection of NAHR of repeat elements in HG002. The mean read length across the two datasets was 22,2±7,2 Kb (ONT) and 16,8±3,3 Kb (PacBio) (Supplemental Figure S6A-B). This difference was expected since the PacBio dataset included libraries subjected to fragment size selection prior to sequencing. After mapping the data on GRCh38, the average diploid genome coverage for the ONT dataset was roughly double that of the PacBio dataset (9.9±2.6 ONT vs 4.3±1.0 PacBio). The TE-reX pipeline detected a total of 17,472 NAHR events in the ONT dataset and 15,521 in the PacBio dataset, with respective average counts of 1,247±370 and 1,336±581 NAHR per library (Supplemental Figure S6C).

**Supplemental Figure S6.**
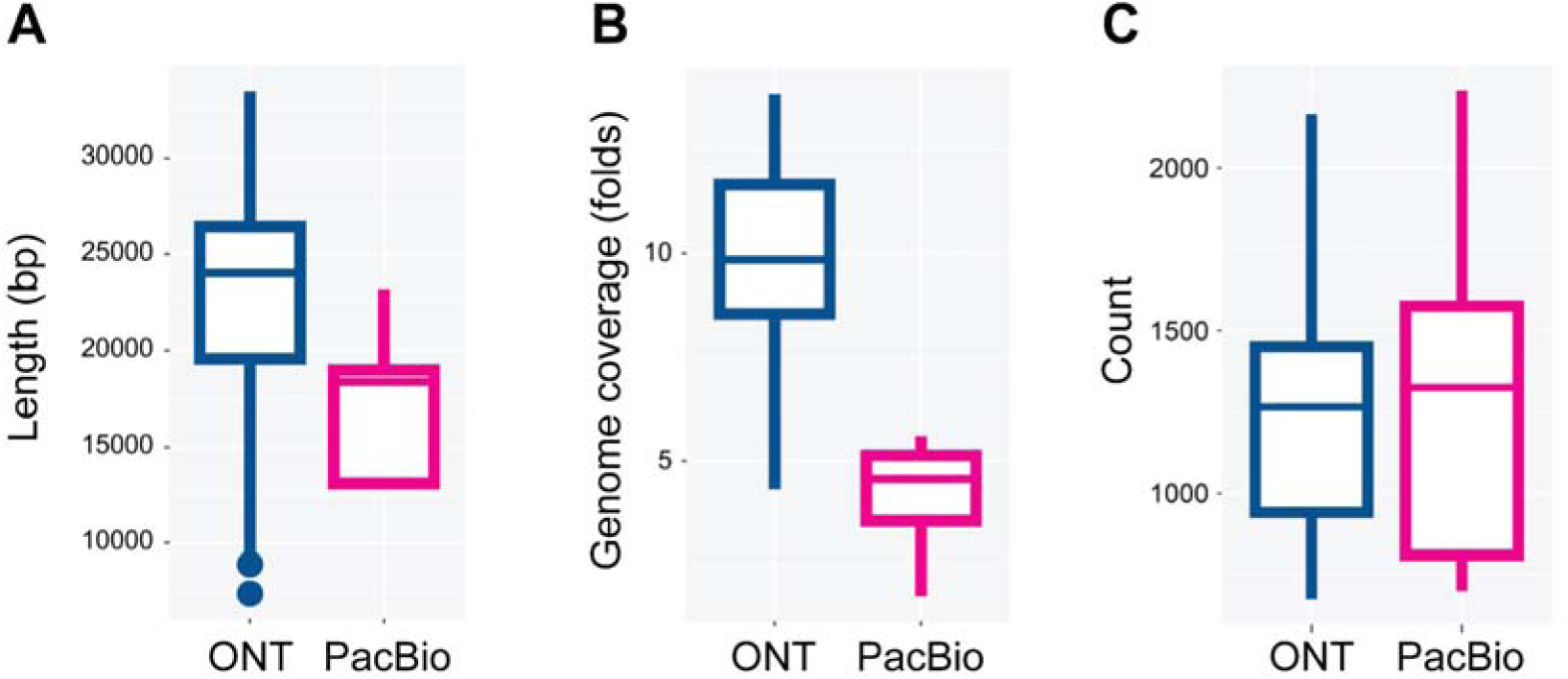
Boxplots showing the distribution of read length (A), diploid genome coverage (B) and total raw NAHR counts (C) in the ONT and PacBio HG002 WGS datasets. The horizontal line in the box shows the median, the box indicates the interquartile range (IQR), whiskers show minimum and maximum values, and dots show outliers with values greater than ±1.5 × IQR.

We expected a number of NAHR events to be detected in both ONT and PacBio dataset, representing polymorphic NAHR or NAHR events generated in the germline of the donor from whom HG002 was established. The main prerequisite for these recurrent NAHR events is that they should be detected in the exact same configuration (i.e., the same pair of repeat elements joined at the same genomic breakpoint) in more than one library. We detected 529 NAHR events in at least two libraries (i.e., one ONT and one PacBio library), with 106 NAHR events found in all 44 libraries. These events were supported by high read counts, showing a high correlation with the count of libraries in which they were detected (r = 0.79, p-value = 2.077e-13; Supplemental Figure S7A). Given the possibility that the ONT and PacBio datasets were obtained from different HG002 samples, we also anticipated that some NAHR events would be detected in multiple libraries within one dataset but not in any library of the other dataset. This platform-specific class of recurrent NAHR could potentially reflect clonal expansions that are differentially represented in the two HG002 source samples. We identified 500 such recurrent NAHR events in the ONT dataset and 1,369 in the PacBio dataset. A vast majority of these platform-specific NAHR events was detected in a few libraries, suggesting a lower copy number compared with recurrent NAHR events found in both datasets. Only 1 NAHR event was detected in 21 ONT libraries and 11 NAHR events were detected in all 22 PacBio libraries (Supplemental Figure 7B-C).

**Supplemental Figure S7.**
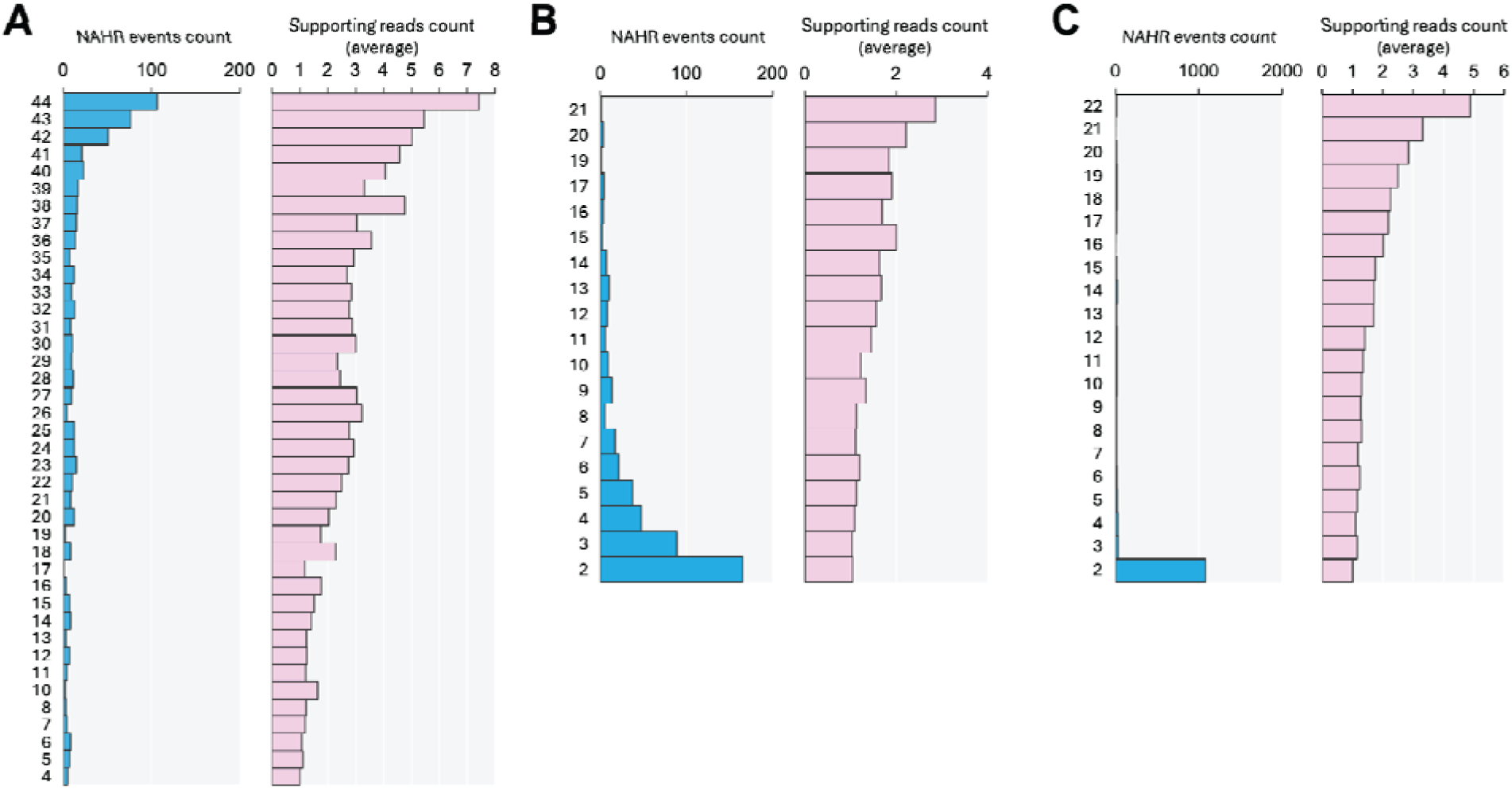
Counts for recurrent NAHR events in the ONT and PacBiodatasets. A) Recurrent NAHR events detected in both ONT and PacBio datasets (left panel; at least in n=1 ONT and n=1 PacBio library) an associated average supporting reads (right panel). B) Recurrent NAHR events detected only in the ONT dataset (left panel; at least in n=2 libraries) and associated average supporting reads (right panel). C) Recurrent NAHR events detected only in the Pacbio dataset (left panel; at least in n=2 libraries) and associated average supporting reads (right panel). The numbers on the y axis of each left panel (blue bars) represent the number of libraries.

Moreover, a comparison of genome-wide profiles of the shared polymorphic/germline and the platform-specific recurrent NAHR events showed relevant differences. While the shared NAHR events and the ONT-only NAHR events were mostly intra-chromosomal, a majority of PacBio-only NAHR events were inter-chromosomal (Supplemental Figure S8A). The shared NAHR and ONT-only NAHR events had the most recombined repeat pairs, displaying a genomic distance of less than 100Kb. In contrast, most intra-chromosomal recurrently recombined repeat pairs in the PacBio-only dataset were distanced more than 100Kb (Supplemental Figure S8B). Most recurrent NAHR events involving proximal intra-chromosomal repeat pairs in the shared and ONT-only NAHR events datasets generated deletions, while inversions were predominant in the PacBio-only dataset (Supplemental Figure S8C); the proportion of SVs generated by repeat pairs distanced >100Kb, and the proportion of repeat families involved was otherwise similar in the three groups (Supplemental Figure S8D-E).

**Supplemental Figure S8.**
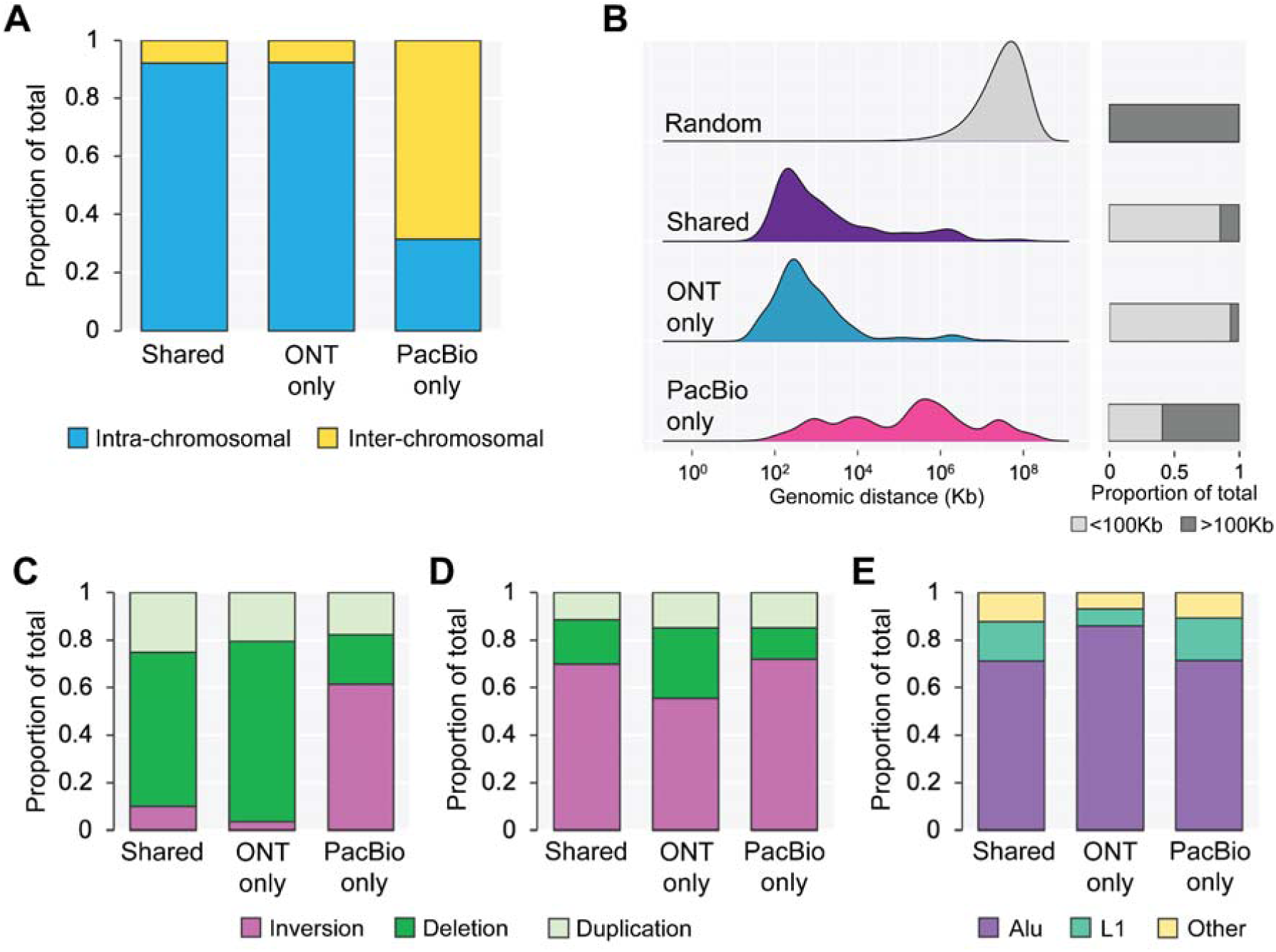
Analysis of NAHR in HG002 WGS libraries sequenced by ONT and PacBio show different genome-wide recombination profiles. **A, B)** NAHR count normalized by sequencing depth (A) and estimates of NAHR per cell (B) in ONT and PacBi HG002 datasets. **C)** Relative proportion of intra- and inter-chromosomal NAHR. **D)** Genomic distance of repeat pairs involved in intra-chromosomal NAHR. The barplots on the right-hand side show the relative proportion of recombined proximal (d<100Kb) and distal (d>100Kb) repeat pairs. **E, F)** Relative proportion of SV types generated by NAHR, for proximal (E) and distal (F) repeat pairs. **G)** Relative proportion of repeat families involved in NAHR. The horizontal line in the box plots shows the median, the box indicates the interquartile range (IQR), whiskers show minimum and maximum values, and dots show outliers with values greater than ±1.5 × IQR.

Subtracting recurrent NAHR events from each dataset produced a list of library-specific, unique NAHR events mostly supported by a single split read (>99.8%). After normalization by sequencing depth, we found that the PacBio dataset had higher counts of unique NAHR events. Similarly, estimates of NAHR per single diploid cell showed higher counts in the PacBio dataset (Figure 3A-B).

**Figure 3.**
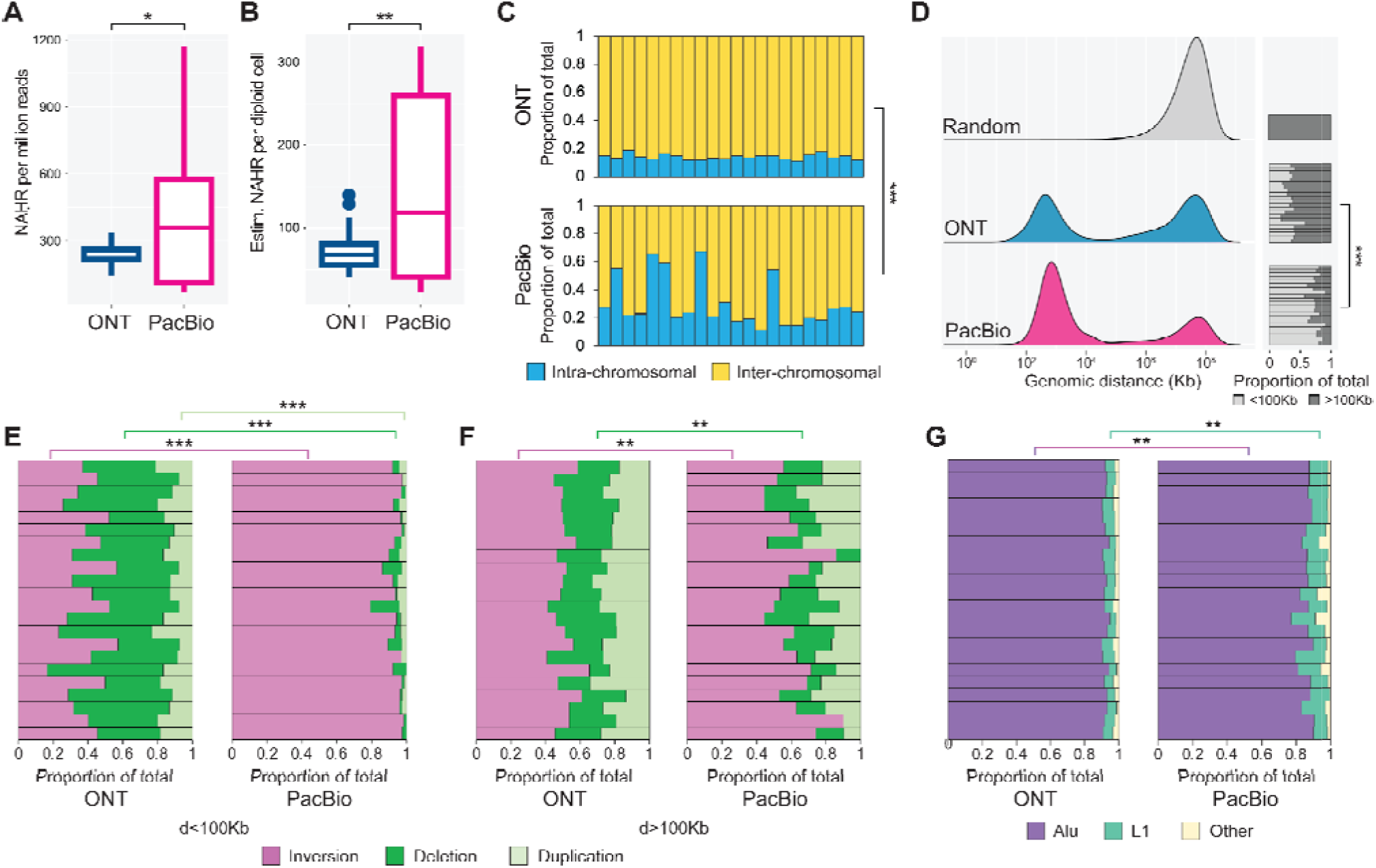
Analysis of NAHR in HG002 WGS libraries sequenced by ONT and PacBio shows different genome-wide recombination profiles. **A, B)** NAHR count normalized by sequencing depth (a) and estimates of NAHR per cell (b) in ONT and PacBio HG002 datasets. **C)** Relative proportion of intra- and inter-chromosomal NAHR. **D)** Genomic distance of repeat pairs involved in intra-chromosomal NAHR. The barplots on the right-hand side show the relative proportion of recombined proximal and distal repeat pairs. **E, F)** Relative proportion of SV types generated by NAHR, for proximal (E) and distal (F) repeat pairs. **G)** Relative proportion of repeat families involved in NAHR. The horizontal line in the box plots shows the median, the box indicates the interquartile range (IQR), whiskers show minimum and maximum values, and dots show outliers with values greater than ±1.5 × IQR. ∗p < 0.05; ^∗∗^p < 0.01; ^∗∗∗^p < 0.001 (Student’s T-test). For bar plots, each bar represents the results for a single library.

Genome-wide profiling of these unique NAHR events revealed significant differences between the two datasets. While most NAHR events were inter-chromosomal, the PacBio dataset exhibited a higher proportion of intra-chromosomal recombination. Additionally, the PacBio dataset showed greater variance compared to the ONT dataset, with five libraries displaying intra-chromosomal NAHR rates exceeding 50% (Figure 3C). Furthermore, intra-chromosomal NAHR between proximal repeat elements was more prevalent in the PacBio dataset (Figure 3D). The relative proportions of SV types generated by NAHR of proximal repeat elements also differed markedly between ONT and PacBio, with inversions being predominant in the PacBio libraries (Figure 3E). Differences were also observed in inversion and deletion rates for NAHR involving distal repeat pairs (Figure 3F). Moreover, the two datasets exhibited varying proportions of repeat families involved in NAHR, with PacBio showing a higher rate of recombined L1 elements and fewer recombined Alu elements compared to ONT (Figure 3G). In summary, these PCR-free WGS datasets revealed several unexpected and platform-specific differences despite being derived from the same cell line.

Nonetheless, the analysis of NAHR in the ONT and PacBio HG002 datasets revealed averages of 83 and 174 recombination events per diploid cell, respectively. Since these estimates were derived from PCR-free libraries of a single cell type with multiple technical replicates, we consider them as a refinement of our earlier estimate of up to ∼5 NAHR events per cell, which was obtained from PCR-based short-read libraries generated from bulk tissues (Pascarella et al. 2022). These findings confirm an important contribution of NAHR of repeat elements to somatic genomic diversity, and validating these data in single cells could help establish NAHR as the most important source of somatic SVs identified to date. However, the study of NAHR in single cells is hindered by the requirement to amplify the tiny amount of genomic material isolated from each cell prior to sequencing. Considering the biases introduced by PCR in NAHR profiles, an ideal single-cell experimental protocol should employ a bias-free and artifacts-free amplification method, which is not yet available. Additionally, such an approach would have to preferentially employ long reads sequencing to maximize the mappability in problematic genomic regions such as segmental duplications, repeat clusters, and centromeres. Only two protocols exist that perform single-cell whole-genome sequencing (scWGS) with long-read technologies, albeit both include PCR (Fan et al. 2021; Xie et al. 2022; Hård et al. 2023). Of these two, SMOOTH-seq has been developed using HG002. SMOOTH-seq can annotate SV in human genomes assembled de novo from scWGS libraries with reads sized at ∼6Kb. To benchmark their technology, the authors of SMOOTH-seq produced libraries from hundreds of single HG002 cells, sequenced by PacBio in the first version of the protocol (Fan et al. 2021) and by ONT in an updated version (Xie et al. 2022). Given the availability of the raw data, we took the chance to compare NAHR in the bulk PCR-free, long reads HG002 datasets and in the scWGS SMOOTH-seq datasets. With this comparison, we aimed at: a) understanding the feasibility of annotating NAHR in state-of-the-art scWGS libraries; b) assessing the impact of PCR on genome-wide NAHR profiles in scWGS libraries; c) verifying whether the differences of NAHR profiles described between ONT and PacBio HG002 bulk datasets are also detected in scWGS SMOOTH-seq datasets. We obtained 157 SMOOTH-seq HG002 libraries sequenced by PacBio and 219 libraries sequenced by ONT. According to the technical details provided by the authors, the differences between the first and second versions of the protocol are small, as they mainly consist of adapting the libraries for sequencing on the ONT platform. Importantly, the number of PCR cycles does not differ greatly, with n=22 for the first version and n=20-22 for the second version. Several SMOOTH-seq ONT libraries were sequenced at higher coverage, but since in our analyses their NAHR profiles did not differ significantly from the remainder, we analyzed the 219 SMOOTH-seq ONT libraries as a whole. After aligning all data on GRCh38 and annotating NAHR events with TE-reX, we noticed that only ∼3% (799/28714) of all NAHR events that are recurrent in the SMOOTH-seq libraries were found also in the bulk WGS ONT and PacBio HG002 datasets. After removing the recurrent NAHR events from each library, a comparison of unique NAHR in SMOOTH-seq and bulk WGS HG002 datasets showed normalized NAHR counts and estimates per single diploid cell ∼15-30x higher in SMOOTH-seq. As in the case of the bulk datasets, the SMOOTH-seq PacBio libraries displayed higher NAHR counts than the ONT libraries (Figure 4A-B).

**Figure 4.**
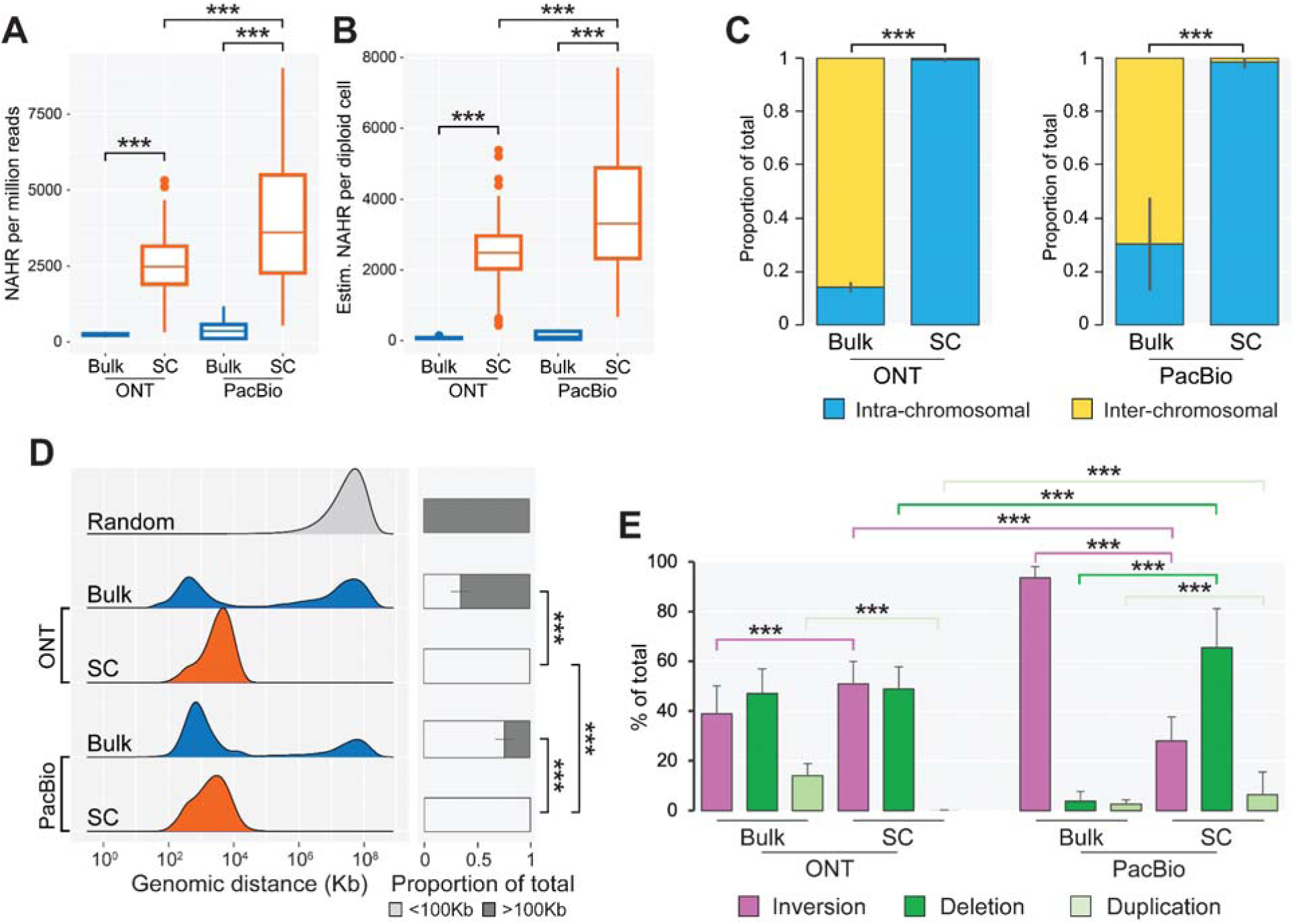
Analysis of recombination events in single-cell SMOOTH-seq HG002 WGS libraries sequenced b ONT and PacBio, and comparison with NAHR in bulk HG002 WGS. **A, B)** NAHR counts normalized by sequencing depth (A) and estimates of NAHR per cell (B) in ONT and PacBio bulk HG002 WGS datasets (“bulk”) compared to recombination events in the respective SMOOTH-seq datasets (“SC”). **C)** Relative proportion of intra- and inter-chromosomal recombination. D) Genomic distance of repeat pairs involved i intra-chromosomal recombination. The barplots on the right-hand side show the relative proportion of recombine proximal (d<100Kb) and distal (d>100Kb) repeat pairs. **E)** Proportion of SV types generated by recombination of proximal repeat pairs. The horizontal line in the box plots shows the median, the box indicates the interquartile range (IQR), whiskers show minimum and maximum values, and dots show outliers with values greater than ±1.5 × IQR. ^∗^p < 0.05; ^∗∗^p < 0.01; ^∗∗∗^p < 0.001 (Student’s T-test). For bar plots, vertical lines represent the standard error of the mean.

Notably, more than 99% of recombination events detected in SMOOTH-seq libraries were intra-chromosomal (Figure 4C) and involving repeat pairs distanced less than 100Kb (median: 3110bp; Figure 4D). Consequently, a characterization of SV generated by NAHR events was possible only for NAHR of proximal repeats, revealing a complex scenario. SMOOTH-seq libraries sequenced by ONT had more inversions and less duplications compared to the respective bulk WGS libraries, whereas SMOOTH-seq libraries sequenced by PacBio had fewer inversions, more deletions, and more duplications with respect to their WGS bulk counterparts. Inversions, deletions, and duplications were also differentially represented in SMOOTH-seq libraries sequenced by ONT or PacBio, with the latter having fewer inversions, more deletions, and more duplications (Figure 4E). Overall, these analyses revealed remarkable quantitative and qualitative differences of NAHR of repeat elements in SMOOTH-seq compared to bulk HG002 WGS libraries and additionally displayed significant cross-platform discrepancies.

## Discussion

We previously showed that somatic recombination of repeat elements happens pervasively in the human genome and we suggested that it may be involved in cell differentiation and in disease. Our findings support the paradigm that structural genomic variants accumulate through life virtually in all tissues and cell types (Martincorena et al. 2015; Loh et al. 2018; Kakiuchi and Ogawa 2021; Grimes et al. 2024). A major challenge in the study of somatic NAHR is the absence of established ground truths or a substantial body of literature to reference when evaluating the quality or significance of new NAHR datasets. Additionally, there are fundamental gaps in our understanding of how NAHR is generated and regulated within the cell. In this context, it is crucial to identify potential technical biases in the annotation of NAHR in NGS data to minimize additional sources of uncertainty. Here, we report that key characteristics of NAHR datasets can be affected by the choice of library preparation methods and sequencing platforms. The results from the capture and sequencing of defined Alu targets demonstrate that, in principle, even a few PCR cycles can generate chimeric molecules indistinguishable from NAHR events. The scenario in real-world libraries is, however, more complex. The analyses of NAHR in WGS short read libraries of the GIAB samples show that low PCR cycling does not generate massive amounts of chimeric recombination artifacts; nonetheless, the differences of NAHR profiles between libraries sequenced by Hiseq or Novaseq are a cause for concern. In general, Hiseq seems to be more resistant than Novaseq to biases introduced by PCR, considering for example that, in Hiseq libraries, the PCR did not affect the proportion of intra-chromosomal NAHR and the distance profiles of recombined repeats. It remains difficult to explain why PCR-free Hiseq libraries have a majority of NAHR events between inverted proximal repeats, while in Novaseq PCR-free libraries, a majority of NAHR involves proximal repeats in direct configuration, although not in all libraries. Assuming comparable sample handling and library preparation protocols for both datasets, possible reasons may lie in the two instruments’ different sequencing chemistry and flow-cell layout, but this hypothesis requires further testing. In light of these results, we recommend against using short-read libraries to study the NAHR of repeat sequences. PCR-free libraries with long DNA reads may seem like a better alternative, offering improved mapping accuracy of genomic fragments containing repeat sequences and less biases. However, profiling NAHR in PCR-free WGS libraries of HG002 has revealed several differences between the ONT and PacBio datasets, which warrant further investigation. It was astonishing that the two datasets exhibited markedly different profiles of intra-chromosomal recombination. Given our limited understanding of the factors affecting NAHR profiles, it is essential to evaluate all potential sources of these discrepancies thoroughly. One possibility is that NAHR events are molecular artifacts generated during library preparation or sequencing. Indeed, molecular artifacts formed by nonspecific DNA ligation have been reported in both ONT and PacBio libraries (White et al., 2017; Griffith et al., 2018). However, chimeric artifacts generated by DNA ligation do not exhibit sequence specificity; they have a random genomic distribution and are highly unlikely to involve the same two DNA fragments more than once in a given library. In contrast, NAHR events annotated by TE-reX feature split read junctions specifically at homologous positions, are not randomly distributed across the genome, and include hundreds of instances where the same two repeats recombine at the same genomic breakpoint. These characteristics rule out the possibility that NAHR events in ONT and PacBio libraries are caused by random DNA ligation. Secondly, differences in NAHR profiles may be due to the varying efficiency of the two platforms in sequencing certain types of SVs. However, to our knowledge, there are no reports of significant qualitative or quantitative differences in SVs discovery in datasets produced by ONT and PacBio that could account for our observations. Finally, the differences in NAHR profiles may be attributed to the use of different HG002 batches in the generation of the ONT and PacBio datasets. This uncertainty regarding the source of the DNA samples allows for speculation; however, even if it is confirmed that the datasets were obtained from distinct HG002 batches, we currently do not know if or how different cell culture conditions (e.g., number of passages, confluency) or variations in library preparation protocols might influence genome-wide NAHR profiles. In summary, we cannot provide a definitive explanation for the observed differences. A pressing reason to address the reliability of NAHR detection in ONT and PacBio data arises from our analyses of NAHR in SMOOTH-seq scWGS libraries. The striking qualitative and quantitative differences in NAHR between bulk and single-cell WGS HG002 datasets suggest that most NAHR events annotated in SMOOTH-seq libraries may be chimeric intra-molecular events generated by PCR. This indicates that current state-of-the-art single-cell WGS approaches are still unsuitable for studying NAHR of repeat elements due to the requirement to amplify the genomic DNA. Therefore, our results show that until a bias- and artifact-free single-cell NAHR assay is developed, long-read PCR-free WGS data remains the method of choice to explore the diversity and impact of NAHR of repeat elements in our genomes.

## Methods

### Genomic DNA purification

Frozen tissues (∼200mg) were reduced to fine powder in a liquid nitrogen-cooled mortar and transferred to 10ml lysis buffer (Laird et al. 1991). Genomic DNA extraction was performed as described previously with minor modifications (Wood 1983). DNA pellets were resuspended in DNA rehydration solution (Promega) and after quantitation with Nanodrop (Thermo Scientific) quality control was performed by running 300ng of resuspended DNA on a 1% agarose gel. The use of human samples in this study was approved by the RIKEN Research Ethics Committee with permission number 2023-15(2).

### Amplification and validation of target Alu sequences

Primers for the 20 Alu targets were designed with Primer3 (Untergasser et al. 2012) and synthesized by Invitrogen. PCR reactions were performed using 25ul of Q5 High-Fidelity Master Mix (NEB), 15ng of genomic DNA, 10uM of forward/reverse primers and H20 in a 50ul total volume. Amplification was carried out with initial 10 cycles of touchdown PCR (1x initial denaturation 98C for 30s, 10x 98C for 10s, primers-specific Tm +10C for 30s decreasing by 1C each cycle, 72C for 7s) followed by 30 cycles of standard PCR (98C for 10s, primers-specific Tm for 30s, 72C for 7s, final elongation 72C for 120s). PCR products were cleaned with AMPure XP beads (Beckman Coulter) and quantified with Qubit High Sensitivity dsDNA assay (Thermo Fisher). Quality control of all amplicons was carried out by running 200ng of cleaned PCR products on a 1.5% agarose gel to verify that a single band was visible for each amplified target. For Sanger sequencing, 20ng of PCR products were transferred in 96-wells plates, mixed with 5uM forward primers and submitted for sequencing to Genewiz.

### Generation of capture-seq libraries

Equimolar amounts of the 20 amplicons were mixed in a 1 to 10 ratio with sheared salmon sperm DNA (Thermo Fisher Scientific) in a final volume of 200ul. Two samples of 50ul volume each were processed for end repair, dA-tailing and sequencing adapter ligation with the NEB Next Ultra II kit (New England Biolabs) according to manufacturer’s instructions with no modifications. The enrichment PCR to introduce the sequencing indexes was carried out for 4 cycles and 8 cycles (“4cyc” and “8cyc”). After purification of the two capture ponds, the capture of Alu sequences, enrichment and post-capture PCR (n=12 cycles) were performed according to our capture-seq protocol previously described with no modifications (Pascarella et al. 2023). The amplified captured fragments were cleaned by AMPure XP beads (Beckman Coulter) and quantified by Qubit High Sensitivity dsDNA assay (Thermo Fisher). The libraries were quality controlled in triplicates with Bioanalyzer High Sensitivity DNA kit (Agilent), pooled in a equimolar fashion, diluted to 4nM and sequenced on a single run of Illumina Miseq using a Miseq Reagent Kit v2 (300 cycles).

### Mapping and annotation of chimeric recombination events in test capture-seq libraries

Overlapping paired reads (median across libraries ∼98%) for the 4cyc and 8cyc libraries were merged in longer contigs using FLASH (Magoč and Salzberg 2011); non-overlapping reads were merged with respective FLASH-extended contigs to generate single FASTQ files that were then mapped on GRCh38 using LAST (Frith and Kawaguchi 2015) as described previously (Pascarella et al. 2023). We used TE-rex to annotate chimeric recombination events in 4cyc and 8cyc libraries and to calculate the identity rate of all recombined amplicon pairs. Command lines and technical details to run TE-reX have been described previously (Pascarella et al. 2023). To distinguish between chimeric molecules generated by template switching and internal priming we intersected the GRCh38 genomic coordinates of each split read available in the TE-reX output with the genomic coordinates of the 20 input Alu sequences, using BEDtools (Quinlan and Hall 2010) with parameters “-wao”. From the intersection output we identified chimeric recombination events that have at least one half of a split read truncated in respect to the original experimental design, indicating internal priming as a mechanism for the chimera formation.

### Annotation of NAHR events in GIAB dataset, WGS bulk HG002 libraries and SMOOTH-seq libraries

Raw fastq files for the different datasets were obtained at sources indicated in the main text. Fastq files were mapped on GRCh38 and NAHR events were annotated with TE-reX as previously described (Pascarella et al. 2023). Post-processing of TE-reX output files was carried out with custom scripts available at https://gitlab.com/mcfrith/te-rex. Estimates of NAHR per diploid cell were calculated by dividing the number of unique NAHR events in a given library by its diploid genome coverage, assuming that 1-fold diploid genome coverage equates to sequencing all the bases in a single diploid cell. We acknowledge that this is an approximation, as coverage is not uniform, with some regions sequenced multiple times and others not at all. Therefore, our estimates may underestimate the true number of NAHR events per diploid cell. However, we consider this an acceptable compromise for now, given the challenges of accurately counting NAHR in single-cell WGS protocols due to PCR artifacts.

### Plotting and statistical analyses

Details about the statistical significance for tests used in this study are described in the Figure legends. Student’s T-test was performed in R. All plots were done in R with package ggplot2 or Excel. Significance of each figure panel is detailed in the respective figure legends. Randomized datasets were generated from the real datasets with varying number of iterations tailored for each specific testing and always comparably sized according to real dataset to avoid any potential bias.

### Data access

All raw and processed sequencing data generated in this study have been submitted to the NCBI Sequence Read Archive (SRA; https://www.ncbi.nlm.nih.gov/sra) under accession number PRJNA1149531. The data can be accessed by the reviewers from this link: https://dataview.ncbi.nlm.nih.gov/object/PRJNA1149531?reviewer=j2r0fi3u7ao2ahgbmp9dbaha3n

## Competing interests statement

The authors declare no competing interest.

## Acknowledgements.

This work was funded by a Research Grant from the Ministry of Education, Culture, Sports, Science and Technology (MEXT), Japan, to the RIKEN Center for Integrative Medical Sciences. P.C. was supported by the RIKEN Aging Project.

## Author contributions

GP devised the project and performed the experiments and data analysis under the supervision of PC and MF. GP, PC and MF wrote the paper.

## Notes

### Competing Interest Statement

The authors have declared no competing interest.

